# The Potential Effect of Vitamin D Supplement on Selected Coagulability Predictors in Vape-Exposed Female Rats

**DOI:** 10.64898/2026.02.25.708056

**Authors:** Aman M. Hammad, Mahmoud Abu Samak, Rana Abu Farha, Lujain F. Alzaghari, Abdelrahim Alqudah, Diana Malaeb, Khaldoun Rasem Shnewer, Souheil Hallit, Muna Barakat

## Abstract

**Background:** Vaping and vitamin D deficiency impact blood coagulation and health. This study aimed to investigate the effects of vitamin D supplementation on coagulation predictors in female rats exposed to E-cigarette vaping.

**Objective:** To examine the effect of vaping alone and vaping with different VD doses on some coagulation predictors, lungs, liver, and kidney functions

**Methods:** Forty-two female Wistar rats were divided into six groups, including vaping and non-vaping with high (50,000 IU) and low (1000 IU) vitamin D doses. Blood samples and histopathological analyses were conducted after one and three months. Nicotine, cotinine, Interleukin-6 (IL-6), D-dimer, coagulation factor X (FX), thrombomodulin (TM), Alanine Transaminase (ALT), and Creatinine levels were analyzed. Additionally, histopathological analyses were conducted on the rats’ liver, kidney, and lung.

**Results:** Exposing rats to vaping for one month caused a significant acute increase in D-dimer, FX, and TM levels to 4402.05 ng/mL ± 785.15, 1.8687 μg/mL ± 0.3132, and 34.71 ng/mL ± 8.42, respectively. However, after three months of exposure, those levels decreased significantly compared to the one-month levels. Supplementation of the vape-exposed rats with a high vitamin D dose reduced levels of IL-6, D-dimer, FX, and TM levels to become 93.285 pg/mL ± 12.715, 439.95 ng/mL ± 294.05, 0.647 μg/mL, and 17.375 ng/mL ± 3.895, respectively, at the end of the three months. Moreover, vaping rats supplemented with the low and high doses of vitamin D had significantly lower nicotine and cotinine levels than the EC group, with a p-value of <0.0001. The histopathological examination revealed that the rat’s lung had necrotic pneumonia when exposed to vaping without vitamin D treatment. Moreover, all vaping groups had an alveolar hemorrhage. Bacterial pneumonia was seen in the high-dose vitamin D vape-exposed group. However, the histopathological examination of the liver indicated no major differences between the groups.

One month of vaping raised D-dimer, FX, and TM levels, which decreased after three months. High-dose vitamin D supplementation reduced IL-6, D-dimer, and FX levels while increasing TM levels after three months. Vaping rats receiving vitamin D had lower nicotine and cotinine levels. Histopathological findings showed necrotic pneumonia and alveolar hemorrhage in vaping rats, with bacterial pneumonia in the high-dose group.

**Conclusion:** Vaping activates inflammatory and coagulation pathways, while high-dose vitamin D appears to mitigate inflammation and blood coagulation issues associated with vaping, potentially aiding in reducing nicotine dependence.

## Introduction

Smoking significantly contributes to early mortality and public health issues globally. E-cigarettes (E-cigs), emerging around 2006-2009, have gained popularity, particularly among young adults, with over 683,300 users in Jordan as of 2020 [1–4].

E-cigs are devices that heat a solution usually composed of propylene glycol or glycerin, nicotine, and flavoring ingredients to produce an aerosol, also known as vapor [5]. While traditional smokers are switching to vaping as a quitting method, concerns arise regarding their role in cardiovascular diseases. Yet, some recent studies concluded that e-cigs are not blameless for causing cardiovascular diseases, including blood coagulation problems. A recent study has shown that the thermal decomposition of vape components generates harmful compounds, which are associated with a significant enhancement in platelet aggregation in vape-exposed mice, suggesting an elevated risk of thrombosis-related cardiovascular disorders[6–8]. Concurrently, vitamin D (VD) is highlighted for its health-modulating effects, including its anticoagulant properties and association with reduced thrombosis risk [9]. Supporting this, a five-year cross-sectional study reported that higher serum levels of VD were significantly associated with a reduced risk of deep vein thrombosis and other serious health outcomes [10].

Based on published literature, this study will focus on female rats due to the higher reported risk of thrombosis in females and the increasing use of e-cigs among women, especially during pregnancy [11]. This research aims to investigate the effects of vape exposure on blood coagulation and inflammation, as previous studies have reported conflicting results. In addition, the study will evaluate the potential protective effect of vitamin D against vaping-induced changes in coagulation and inflammatory markers.

## Materials and methods

### Study design

VOOPOO DRAG M100S vape device with PnP-TM2 0.8 Ohms coil resistance, VGOD berry bomb (sour strawberry belt) flavor, 50% propylene glycol, 50% vegetable glycerin, and 18mg/ml nicotine. Hi Dee^®^ (2000 IU VD/5 drops) and tera D^®^ (400 IU VD/ml) were used. The study was conducted in two phases as follows:

### Phase one

Three groups of female rats were exposed to vaping for one month, and the other three groups were not exposed to vaping nor received any treatment.

### Phase two

After the first month of exposure, VD was given to two vaping and two non-vaping groups for 2 months. The two vaping groups received VD at different dosages—one with a low dose (1000) and the other with a high dose (50,000 IU weekly). Additionally, two other rat groups will receive vitamin D without vape exposure, one with a high dose (50,000) and the other with a low dose (1000 IU daily).

### Animal management

This study was an in vivo study which used 42 Wistar rats, aged 12 weeks, housed at Applied Science Private University in Amman, Jordan, for acclimatization over one week under standard laboratory conditions, including a temperature of 21–23 °C, relative humidity of 35–70%, and a 12-hour light/dark cycle, with free access to food and water.

All experimental procedures involving animals were conducted in accordance with institutional and international guidelines for the care and use of laboratory animals. Ethical approval was granted by the Research and Ethics Committee of the Faculty of Pharmacy, ASU (Approval No.: 2024-PHA-40).

Rats were randomly assigned to six experimental groups (n = 7 per group) as follows: Control group (C): This group was not exposed to vaping nor VD treatment for the entire 12-week period (negative control). Vaping group (EC): This group was exposed to vaping only for the entire 12 weeks, 2 hours per day, 5 days a week, without receiving VD treatment (positive control). Vaping + low dose VD (ECD1000): This group was exposed to vaping for 4 weeks. Then, 1000 IU of VD was administered daily for 8 weeks while continuing vaping exposure. Vaping + high dose VD (ECD50,000): This group was exposed to vaping for 4 weeks. Then, 50,000 IU of VD was administered once weekly for 8 weeks while continuing vaping exposure. Low-dose VD (D1000): This group took 1000 IU of VD daily for 8 weeks without exposure to vaping. High dose VD (D50,000): This group took 50,000 IU of VD weekly for 8 weeks without exposure to vaping.

### The exposure settings

The vaping groups were removed from their cages to a 50 cm (length) x 50 cm (width) smoking chamber shown in **Figure.1**, and back again to their cages after the end of the exposure. Rats were exposed to the vape twice daily in two separate sessions, one hour in the morning and one hour in the evening. After modification, an air pump was used to pull the vapor from the mouthpiece through plastic oxygen tubes directly connected to the smoking chamber. At the beginning, the chamber was filled with vapor (saturation phase). In the saturation phase, the pump was activated for 20 seconds. During this time, the fire button was pressed for 5 seconds, followed by 2-3 seconds of rest to avoid coil and mouthpiece overheating. The second exposure phase began after the chamber was filled with vapor. During this phase, the pump was activated for 5 seconds, followed by 20 seconds of rest, mimicking real situations of typical user behavior.

After the first hour of exposure, the pump was stopped, and the chamber remained closed until the rats inhaled the remaining vapor inside. Then, the chamber was opened and supplied with fresh air by opening a window directly beside the chamber, and the rats remained inside until the next hour of exposure. The exact process was repeated in the second hour of exposure.

### VD treatment

VD was given orally by directly dropping the dose into the rat’s mouth after one month of vape exposure for 4 groups. It was administered in two doses: 1,000 IU daily and 50,000 IU weekly. The equivalent doses were calculated using this equation: animal equation dose = Human dose/60 × Km ratio, Km = 6.2 [12]. The equivalent dose of 50,000 IU was 5 drops from the commercially available Hi Dee® (2000 IU/5 drops) vial and 3 drops from the commercially available tera D® (400 IU/ml) vial for a 1000 IU dose. The vitamin was given in the morning.

### Blood sampling

Blood samples were collected in ethylenediamine tetraacetic acid (EDTA) tubes. The blood plasma was obtained and stored in Eppendorf tubes at -80 °C.

### Measurement of nicotine and cotinine levels

Nicotine and cotinine concentrations were measured using LC-MS-8030 As LIQUID CHROMATOGRAPH MASS SPECTROMETER-Triple Quad MS (Shimadzu Corp.Japan). Briefly, a C18 4.5 mmx15cm x 0.2um column was used as a stationary phase, with a mobile phase of 75% acetonitrile mixed with 0.05% formic acid and 25% distilled water, eluted isocratically at a flow rate of 0.3 ml/min for 4 minutes. The MS interface was electrospray ionization (ESI) running in positive ionization mode to generate [M+H]+ ions at m/z 162.23, 176.21 for nicotine and cotinine, respectively.

### Measurement of IL-6, D-dimer, TM, and FX levels

Commercially available rat-specific ELISA kits (ELK Biotechnology, Wuhan, China) were used to detect IL-6, D-dimer, and TM, according to the manufacturer’s instructions. Briefly, Plasma samples, standards, and controls were placed in wells pre-coated with particular capture antibodies, then incubated and washed. The detection antibodies were then coupled with horseradish peroxidase (HRP), and colorimetric detection was performed on a TMB substrate. The reaction ended with a sulfuric acid solution, and the absorbance was measured at 450 nm using a microplate reader. The concentrations were determined using standard curves developed from the established values included in each kit. Each sample was examined in duplicate.

FX levels were evaluated using a mouse-specific sandwich ELISA kit (Mouse F10 ELISA Kit, Reed Biotech Ltd., Hubei, China), following the manufacturer’s protocol, which was similar to the previously used method. Although the kit was designed for mouse samples, it was used due to documented cross-reactivity with rat plasma. Each sample was examined in duplicate.

### Measurement of ALT and Creatinine levels

Serum alanine aminotransferase (ALT) and creatinine levels were measured using the BioSystems ALT and Creatinine kits (BioSystems S.A., Barcelona, Spain). ALT activity was assessed by monitoring NADH oxidation at 340 nm via spectrophotometry, while creatinine levels were determined through the Jaffé reaction, which forms a colored complex measured at 500 nm.

### Histopathological analysis

Lung, kidneys, and liver were harvested from three rats from each group after sacrifice. Organs were fixed in 10% formalin for 1 week. Then, the organs were transferred to Smart Lab to make the histopathological analysis. The organs were trimmed into cassettes and then a high concentration of alcohol was added to dehydrate the tissues. A clearing agent (xylene) was added to replace the alcohol. Then, melted paraffin wax was added to support the tissues. Using a microtome, the paraffin block was cut into 4-µm-thick sections and placed on a glass slide to be stained with hematoxylin and eosin for examination under the light microscope.

### Statistical analysis

All results were expressed as mean ± standard deviation. Data were analyzed using the two-factor analysis of variance (ANOVA) in GraphPad PRISM, the tenth version of statistical software to calculate the statistical significance between the groups at different times. Tukey’s post hoc test was then used to compare the differences between the groups, considering a P value of <0.05 a statistically significant value. In addition, the Pearson test was used to detect the correlation between nicotine concentration and other parameters in the vaping groups.

## Results

### Nicotine and Cotinine

As shown in **Fig. 2**, after the first month of exposure, the nicotine and cotinine levels increased significantly from 0 ng/mL to 5.5 ng/mL ± 0.1 and 103 ng/mL ± 5, respectively. After 3 months, a significant increase occurred in both compounds’ mean levels to 12.45 ng/mL ± 0.55 in nicotine and 283.5 ng/mL ± 3.5 in cotinine, with a p-value of <0.0001 in the EC group.

**Figure 1.**
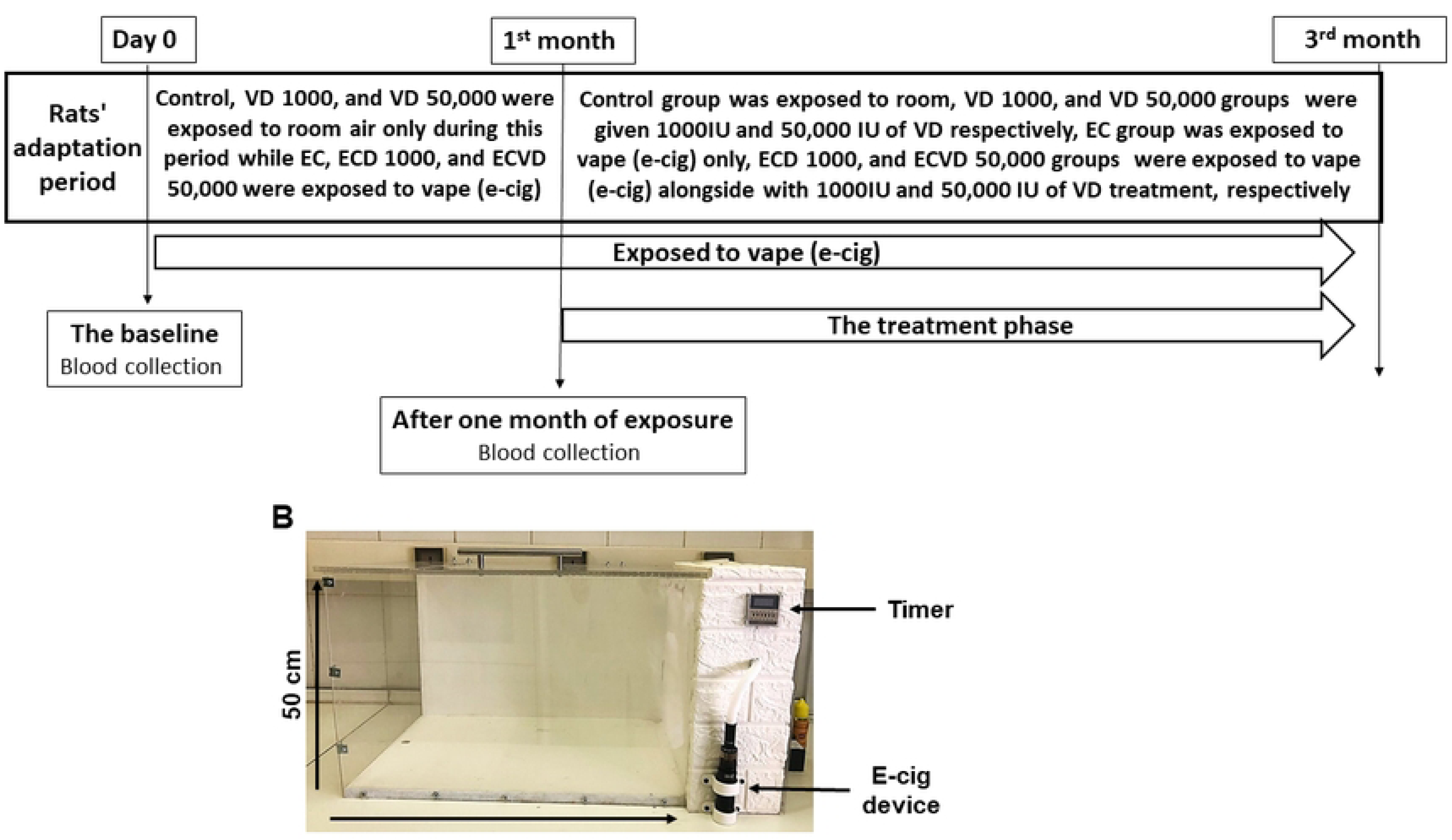
(A) The timeline of the experiment, and (B) the vaping chamber.

**Figure 2.**
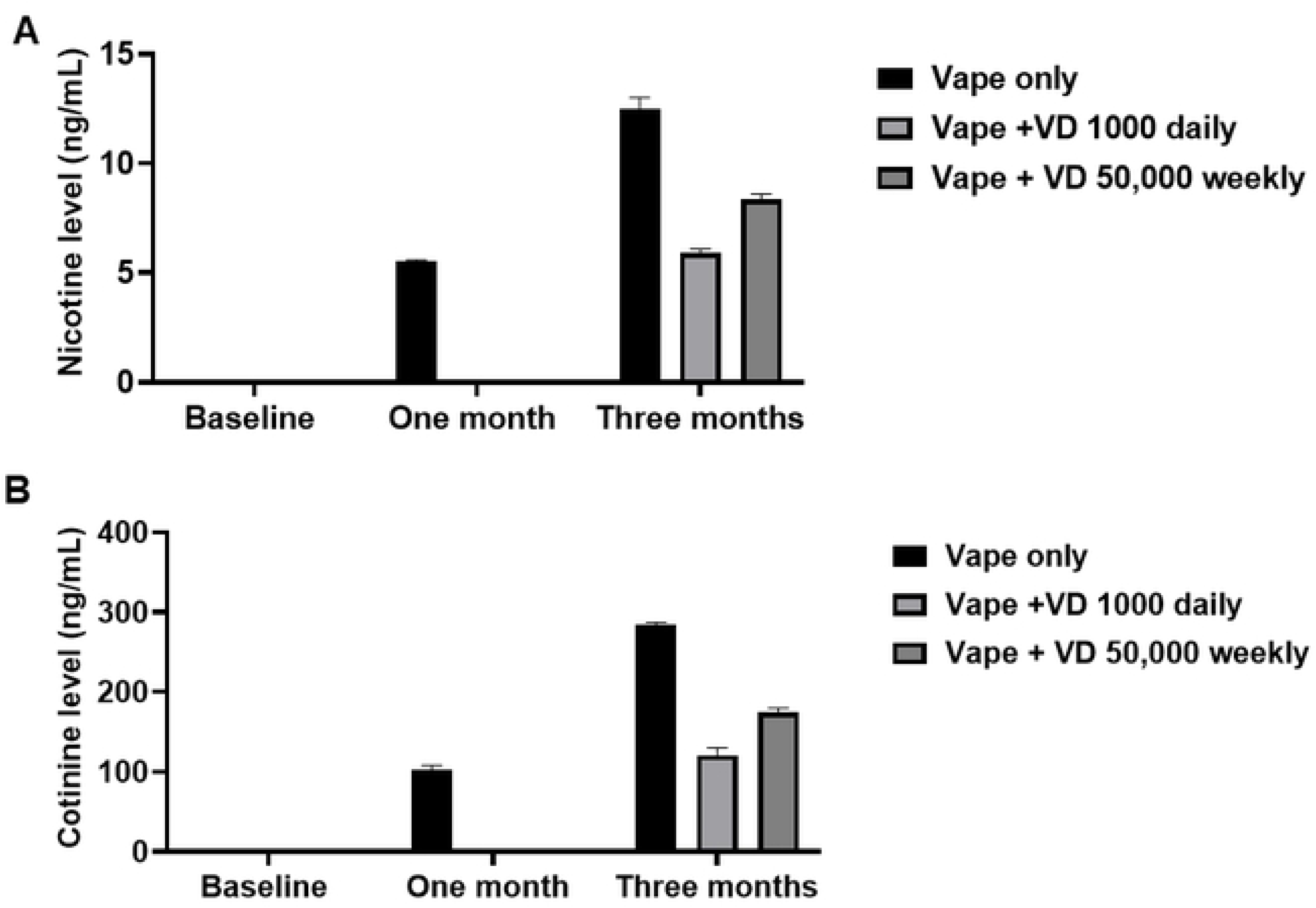
The (A) nicotine and (B) cotinine concentrations (ng/mL) among all vaping groups at baseline, one month, and two months.

However, the ECD1000 group had nicotine concentration close to the one-month level, with a slight increase of 7.27%. Consequently, the cotinine levels increased significantly in three months to 120 ng/mL ± 9.5, with a p-value of 0.0010.

Regarding the ECD50,000 group, the nicotine mean level was 8.25 ng/mL ± 0.25, and the cotinine mean level was 174 ng/mL ± 6, representing a statistically significant increase in nicotine and cotinine levels after three months of exposure to a p-value of <0.0001. Pearson test results indicated that there was no correlation between nicotine level and other parameters.

### Interleukin-6 (IL-6)

At baseline, the mean IL-6 level was 71.843 pg/mL ±10.414. After 1 month, the IL-6 level was 97.19 pg/mL ± 14.4294, representing a slight, non-significant increase of approximately 35%. After 3 months, no statistically significant difference was observed in the EC group. The mean IL-6 level was 115.1 pg/mL ± 10.6, representing 60% and 18% higher levels than the baseline and one-month levels, respectively.

For the VD-treated groups, the mean value for the D1000 group was 108.175 pg/mL ± 28.325, which was 1.5-fold higher than the baseline level. The mean value for the D50,000 group was 235.9 pg/mL ± 109.1, which was significantly higher than both the D1000 group and the C group, with a p-value of <0.0001.

Among the vape-exposed groups, the lowest IL-6 levels were in the ECD50,000 group, with a mean of 93.285 pg/mL ± 12.715, and the highest levels were in the ECD1000 group, with a mean of 122.23 pg/mL ± 46.67. Group ECD1000 had approximately 31% higher, non-significant IL-6 levels compared to group ECD50,000.

As **Figure 3** summarizes, group D50,000 had significantly higher IL-6 mean levels than the EC, ECD1000, ECD50,000, and C groups, with p-values of 0.0002, 0.0004, <0.0001, and <0.0001, respectively. Group ECD1000 had approximately 13% higher IL-6 than the D1000 group.

**Figure 3.**
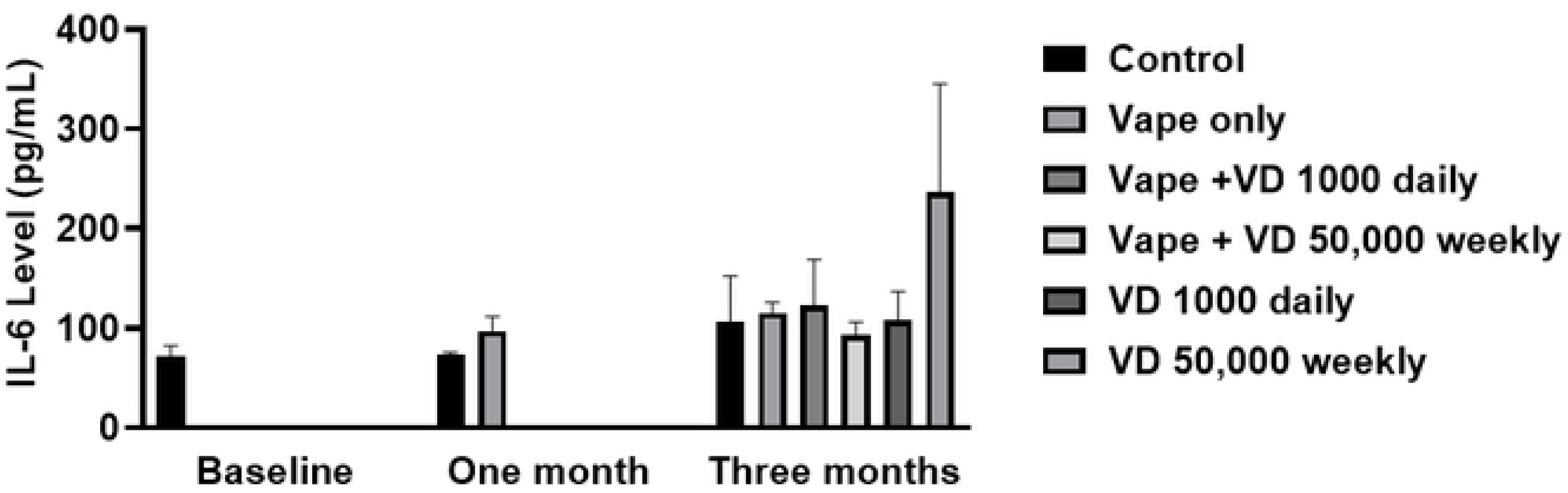
Summary of mean plasma IL-6 concentrations (pg/mL) among all the groups at baseline, one month, and three months.

### D-dimer, FX, and TM

At baseline, the mean D-dimer, FX, and TM levels were 1412.13 ng/mL ± 1072.1288, 0.8027 μg/mL ± 0.0945, and 7.606 ng/mL ± 1.6511, respectively. After 1 month, all the parameters increased significantly after the first month of exposure. The D-dimer levels increased to 4402.05 ng/mL ± 785.15, with a p-value of 0.0019. Moreover, the FX level increased to 1.8687 μg/mL ± 0.3132 with a p-value of < 0.0001, and the TM level increased to 34.71 ng/mL ± 8.42 with a p-value of <0.0001. After 3 months, in the EC group, the mean D-dimer level decreased significantly with a p-value of 0.0016 from the one-month level to become higher by only 3% than the C group reading, with mean values of 1370.65 ng/mL ± 853.05 and 1332.25 ng/mL ± 376.85, respectively. The FX level was 0.904 µg/ml ±0.233, which was lower by 106.71% than the one-month level. The mean level did not differ significantly from the C group, which had a mean value of 0.9975 µg/ml ± 0.3965. The mean level of TM was 13.475 ng/mL ± 2.125, which represented a statistically significant decrease with a p-value of <0.0001. This TM mean value is lower than in the C group by 5.79%, indicating a non-significant difference.

After VD treatment, D-dimer levels did not differ significantly between the VD-supplemented groups. Group D1000 had the highest D-dimer level compared to group D50,000 and the C group, with mean values of 2345.95 ng/ml ± 1585.65, 1785.05 ng/ml ± 1420.05, and 1332.25 ng/ml ± 376.85, respectively. Group D1000 had approximately 31.42% higher levels than group D50,000 and 76.07% than the C group. In group D50,000, the d-dimer level was higher by 34% than in the C group.

The mean D-dimer level in the ECD1000 group was 956.9 ng/mL ± 589.6, which decreased significantly from the one-month mean level with a p-value of 0.0003. The ECD50,000 group had a D-dimer mean level of 439.95 ng/mL ± 294.05, representing the lowest D-dimer level compared with the EC and ECD1000 groups. Although the difference between the group’s levels was not statistically significant, group ECD50,000 had 117.5% and 211.6% lower levels than the ECD1000 and EC groups, respectively. Both VD supplemented with vape exposure groups had lower D-dimer levels than the C group. The C group had approximately 39.23% higher levels than group ECD1000 and 202.8% than group ECD50,000.

The analysis showed no statistically significant differences between the groups, as shown in **Figure S4A**. However, the D-dimer levels in the D1000 and D50,000 groups were higher than those in the ECD1000 and ECD50,000 groups by 145.17% and 305.78%, respectively. The lowest D-dimer levels were found in group ECD50,000 among all other groups.

The highest increase in FX mean level among all the groups was found in group D50,000, which had a significant elevation from the baseline level to 1.5575 μg/mL ± 0.2335 and a p-value of <0.0001. A statistically significant difference between group D50,000 from group D1000 mean value of 0.9275 μg/mL ± 0.0235, and the C group mean value of 0.9975 μg/mL ± 0.3965, appeared after analysis with p-values of 0.0015 and 0.0062, respectively. In contrast, group D1000 had a higher FX mean level of 15.56% from the baseline and a lower mean level of 7.55% from the C group.

The FX mean level decreased significantly from the one-month mean value to 0.752 μg/mL ± 0.245 with a p-value of <0.0001 in the ECD1000 group. a non-statistically significant decrease in FX mean level by 20.21% in the ECD1000 group compared to the EC group. The lowest FX mean level was found in group ECD50,000, which was equal to 0.647 μg/mL. Even not statistically significant, the ECD50,000 group had a 16.23% lower mean FX level than the ECD1000 and 39.72% than the EC groups. Both the ECD1000 and the ECD50,000 groups had lower FX levels than the C group by 32.64% and 54.17% respectively.

According to **Figure S4C**, group D50,000 had a significantly higher mean FX level than all the groups. When group D50,000 was compared with vaping groups, the results were different p-values of 0.0009 against the EC group, <0.0001 against group ECD1000, and <0.0001 against group ECD50,000. The C group had the second-highest mean FX level, followed by group D1000 and the EC group, which had close mean values. Group D1000 had a higher mean FX level of 43.36% compared to group ECD50,000 and 23.34% compared to group ECD1000. However, group D1000 had a lower FX mean value than the EC group by 2.60%.

Regarding TM, a slight, non-significant decrease of 23.95% compared to the baseline level was detected in group D1000, which had a mean value of 6.1365 ng/ml ± 2.4315. In contrast, group D50,000 showed an increase in TM level by 5.47% from baseline with a mean value of 8.022 ng/ml ± 1.251. Group D1000 had a lower TM mean value of 30.73% than group D50,000. The D1000 and D50,000 groups had no statistically significantly lower TM mean values than the C group, which had a mean level of 14.255 ng/ml ± 3.17002, by 132.32% and 77.68%, respectively.

In addition, the TM mean level decreased significantly in the ECD1000 and ECD50,000 IU VD groups, with mean values of 20.41 ng/mL ± 8.35 and 17.375 ng/mL ± 3.895, respectively, and corresponding p-values of 0.0002 and <0.0001. The highest TM mean level was in group ECD1000, which was higher than the EC group by 51% and by 17.47% compared to group ECD50,000. Both the ECD1000 and the ECD50,000 groups had higher, but non-significant, TM mean values compared to the C group, by 43.18% and 21.89%, respectively.

When comparing all the groups, the group receiving D1000 had a significantly lower TM mean level than both the ECD1000 and ECD50,000 groups, with p-values of 0.0006 and 0.0106, respectively. Regarding group D1000, a non-statistically significant difference of 119.59% lower TM mean level than the EC group was found after analysis. The D50,000 group had a significantly lower mean value than group ECD1000 IU VD, with a p-value of 0.0037 and a non-significant lower mean level than group ECD50,000 IU VD by 116.64%. In addition, group D50,000 had a lower TM level of 67.99% than the EC group. **Figure S4B** visually summarizes those findings.

### ALT and creatinine

At baseline, the ALT and creatinine mean levels were 60.1 U/L ± 4.1605 and 0.645 µmol/L ± 0.015, respectively. After 1 month, ALT level decreased significantly after one month of exposure to 42.7 U/L ± 3.8105, with a p-value of 0.0028. However, the Creatinine level was 0.69 µmol/L ± 0.09, which was considered a non-significant increase by 6.98% from baseline levels. After 3 months, in the EC group, the ALT mean level increased significantly to 57.5 U/L ± 3.5 and a p-value of 0.0136, less by 4.33% than the baseline and 15.38% than the C group, which had a mean value of 67.95 U/L ± 1.65. A 3% decrease in creatinine levels occurred compared to the first month’s levels to be 0.67 µmol/L. However, compared to the control’s mean level of 0.61 µmol/L ± 0.03, the EC group had 9.84% higher creatinine levels.

After VD treatment in both groups, an increase in ALT levels appeared. Group D1000 had 15.35%, and group D50,000 had 13.71% higher values than the baseline, with mean values of 71 U/L ± 8.5 and 69.65 U/L ± 7.85, respectively. Compared with the C group, group D1000 and group D50,000 had higher mean ALT levels by 4.30% and 2.44%, respectively. Both groups, D1000 and D50,000 had very close mean values, with group D1000 having approximately 1.9% higher values.

Regarding creatinine, group D1000 had the highest creatinine level of 0.665 µmol/L ± 0.005 by 3.10% and 9.02%, respectively, compared with group D50,000 and the C group. When compared with the C group, group D50,000 had a mean level of 0.645 µmol/L ± 0.015, which was 5.74% higher.

Both the ECD1000 and the ECD50,000 group’s ALT mean levels increased compared to one month after VD treatment, with mean values of 75.25 U/L ± 16.15 and 51.9 U/L ± 5.2, respectively. This increase was significant in the ECD1000 group with a p-value of <0.0001, and non-statistically significant in group ECD50,000, with a 21.54% increase only. The ECD1000 group had the highest ALT mean level among the EC group and group ECD50,000, with p-values of 0.0015 and 0.0001, respectively. Group ECD50,000 had a 9.74% lower ALT mean value than the EC group.

**Figure S5A** showed that the highest ALT mean level was in group ECD1000 compared to all other groups. It had a significantly higher mean level of 75.25 U/L ± 16.15 than the EC group with a p-value of 0.0051. Both groups D1000 and D50,000 had lower ALT mean values of 5.65% and 7.44% than the ECD1000, respectively. In addition, group ECD1000 had a 9.70% higher mean ALT level than the C group.

However, the group ECD50,000 mean ALT level was 9.74% lower than the EC group. Upon analysis, the ECD50,000 group had a significantly lower ALT mean value than the D1000, D50,000, and the C groups, with p-values of 0.0021, 0.0051, and 0.0145, respectively.

The creatinine in the ECD1000 group was significantly higher than the EC group and the C groups, with a mean value of 0.78 μmol/L ± 0.01 and p-values of 0.0120 and <0.0001, respectively. In contrast, the ECD50,000 group had a mean creatinine level of 0.72 μmol/L ± 0.06, which was 7.69% less than the group ECD1000 IU VD mean level and higher than the EC group by 7.46%. A significantly higher mean creatinine level with a p-value of 0.0120 was found between the ECD50,000 and the C group.

A significantly higher mean creatinine level was observed in group ECD1000 than in the D1000 and D50,000 groups, with p-values of 0.0076 and 0.0011, respectively. Regarding group ECD50,000, it had an 8.27% higher mean creatinine level than group D1000 and 11.635% higher than group D50,000. There were very close mean values in the EC, D1000, and D50,000 groups. The EC group had 0.75% higher creatinine levels than group D1000 and 3.88% higher than group D50,000, as illustrated in **Figure S5B.**

### Histopathological examination

#### Liver

The EC and the C groups had a resemble hepato-microscopic examination with moderate lymphocytic infiltration in the portal tracts, and no significant fibrosis, steatosis, or cirrhosis was observed, as shown in **Figure S6A and S6B.**

The hepatic tissue in group D1000 resembled the C group by having mild chronic inflammation in the portal tracts, with a predominantly lymphocytic infiltrate without significant hepatocellular damage or fibrosis. As shown in **Figure S6C**. In contrast, group D50,000 had normal liver tissue without significant fibrosis or inflammation. As shown in **Figure S6D**.

Histopathological examination reveals chronic inflammation with lymphocytic infiltration and occasional macrophages in the portal tracts in the ECD1000 group, without signs of fibrosis, necrosis, or steatosis, see **Figure S6E**. Moreover, in the ECD50,000 group, the liver showed evidence of chronic inflammation with mild lymphocytic infiltration around the portal tracts without significant hepatocellular damage or fibrosis, see **Figure S6F**.

### Kidneys

In the C group, the glomeruli demonstrate cellular expansion in both mesangial and endocapillary areas, with reduced capillary lumen size. Tubules show preserved morphology or mild nonspecific changes, as illustrated in **Figure S7A**.

The glomeruli in the EC group exhibited diffuse mesangial hypercellularity and mild endocapillary proliferation, with some narrowing of capillary lumina. There was no significant glomerulosclerosis or crescent formation, and the tubules appeared largely preserved, as shown in **Figure S7B**. In group D1000, the glomeruli demonstrate mild mesangial expansion with focal endocapillary hypercellularity. Tubules are largely preserved, with minimal signs of atrophy and rare protein casts, as **Figure S7C** provides.

In group D50,000, Glomeruli showed mesangial and endocapillary hypercellularity with capillary lumen narrowing. There was no evidence of crescent formation or significant glomerular scarring. Tubules are unremarkable or show mild reactive changes, as **Figure S7D** provides.

In the ECD1000 group, as shown in **Figure S7E**, glomeruli showed mesangial and endocapillary hypercellularity with capillary lumen narrowing. Tubules were unremarkable or showed mild reactive changes. Blood vessels were within normal limits, without evidence of vasculitis or thrombosis, which indicates glomerular inflammation or injury.

The glomerular inflammation and injury were more severe in the ECD50,000 group, as shown in **Figure S7F,** which was characterized by the expansion of the mesangial matrix with focal crescent formation and endocapillary hypercellularity.

### Lungs

In the C group, the lung tissues appeared with no significant inflammatory infiltrates, numerous blood cells, and preserved alveolar walls, as represented in **Figure S8A**. However, in the EC group, the alveolar spaces were filled with abundant neutrophils, and areas of necrosis were evident, consistent with necrotizing pneumonia. The alveolar walls exhibit marked disruption and extensive hemorrhage in the affected regions, as shown in **Figure S8B**.

Diffuse interalveolar hemorrhage was evident in group D1000, with blood in the alveolar spaces and mild edema in the interstitium without any alveolar wall necrosis or significant inflammation observation, as evidenced by **Figure S8C**. In contrast, group D50,000 had normal lung tissue without hemorrhage, fibrosis, or inflammation, as evidenced by **Figure S8D**.

Extensive red blood cells were present in the alveolar spaces in the ECD1000 group, indicating hemorrhage. The alveolar walls were intact, but blood-filled spaces were noted, with occasional hemosiderin-laden macrophages. Mild interstitial edema was present without significant inflammation, as illustrated in **Figure S8E**.

In the ECD50,000 group, the lung was more injured, as evidenced in **Figure S8F**, by having the alveolar spaces filled with neutrophils, indicating acute bacterial pneumonia. The alveolar walls showed mild inflammation, and a few alveolar septa were thickened due to edema. The mucopurulent exudate was present within the bronchioles, and the surrounding alveoli were congested.

## Discussion

The extensive advertising of e-cigs as a safer and healthier alternative to traditional tobacco smoking has considerably boosted their popularity, particularly among teenagers and young adults. The appealing tastes and generally moderate odor of vaping devices have particularly attracted female consumers, emphasizing the importance of researching any potential risks linked with their usage. According to a global survey conducted in 2020, an estimated 68 million people are active e-cig users globally [13]. A survey conducted by a team of British scientists found that the majority of vape shop consumers are between the ages of 18 and 25, with females accounting for around 41% of this population. Moreover, information received from vape shop workers suggested that fruit-flavored e-liquids were the most popular option among their customers [14]. According to emerging data from several research, tobacco smoking has a deleterious impact on VD status by interfering with key enzymes involved in its production and metabolism. Furthermore, smoking has been linked to increased liver damage indicators, which may inhibit VD synthesis and lead to a greater risk of deficiencies among tobacco users [15]. This study is particularly significant because there is little information on the consequences of VD on e-cig users, particularly in terms of blood coagulability parameters and inflammatory markers.

The current study’s principal findings show that one month of exposure to e-cig vapor led in a substantial increase in blood levels of nicotine, TM, FX, D-dimer, and IL-6. These findings are consistent with previous studies that found similar short-term effects of vaping on coagulation indicators and inflammatory markers [16, 17]. The significant rise in certain clinical markers might be attributed to nicotine or the vape juice ingredients themselves. Many in vivo and in vitro investigations revealed that nicotine, flavoring ingredients, hygroscopic carriers, and metals emitted by the heated coil might cause cardiac toxicity and increase IL-6 and other inflammatory markers. likewise, previous research found that acute vape exposure raises the risk of cardiovascular disease via worsening endothelial dysfunction [18, 19]. However, during the third month of exposure, the levels of these biomarkers were decreased, except for nicotine in the vape-only group and IL-6, which remained high. Notably, nicotine concentrations were lower in both the vape + 1000 IU VD and vape + 50,000 IU VD groups, indicating a possible modulatory impact of VD supplementation. In support of this, Knihtilä et al. found that appropriate maternal VD levels during pregnancy were related with reduced cotinine concentrations in tobacco-exposed mothers, which contributed to better respiratory outcomes in their children [20]. The study also identified a significant interaction between cotinine and VD levels, though the underlying mechanism remains unclear. This interaction may be attributed to the anti-inflammatory and antioxidant properties of VD [15].

The observed increases in IL-6 might be attributed to the increase in toxic substances generated by vaping and their buildup in the lungs. These irritating substances produce oxidative stress, which increases the production of white blood cells and cytokines. The observed decrease in blood coagulation indicators after one month of exposure might be due to the rats’ physiological ability to adapt and maintain homeostasis over time [21]. This conclusion is supported by Garrett’s study, which examined the long-term effects of cigarette smoke exposure on coagulation in rats. In this study, rats exposed to tobacco smoke for 22 weeks showed no significant variations in plasma clotting times compared to the control group, indicating an adaptive response. Furthermore, the study revealed an age-dependent effect, with the prothrombotic effect of cigarette smoke being more prominent in older rats (24 months) than in younger rats (about 3 months), indicating that age may influence susceptibility to smoking-induced coagulopathy [22]. In research on the cardiovascular impacts of vaping, Dai et al. exposed 6-week-old rats to e-cig aerosol for 5 hours per day, 4 days per week, for 3 months. Their findings showed that this exposure regimen did not cause substantial changes in blood pressure or heart rate at the end of the research [23]. To further explore the temporal dynamics of cardiovascular adaptation to vaping, El-Mahdy et al. conducted a prolonged exposure study in which rats were subjected to e-cig vapor for 60 weeks, with pulse measurements recorded at multiple intervals. During the initial 8 weeks, pulse rates were elevated relative to baseline values, suggesting an acute physiological response. However, from weeks 8 to 16, pulse rates declined, indicative of adaptive mechanisms. Following this period, from week 16 onward, pulse rates increased progressively and significantly, persisting through the remainder of the 60-week exposure period [24]. Additionally, Rafiq et al. confirmed an inverse connection between IL-6 and TM. In individuals with coronary artery disease, NF-kB was down-regulated whereas TM expression was up-regulated. In vitro and in mouse lung injury models, inhibiting NF-kB activity reduced cytokine-induced TM downregulation [25]. These prior findings are consistent with the findings of this investigation, which showed that TM and IL-6 levels were conflicting [26].

Recently, in a vitro study conducted by Cirillo et al, tissue factor expression at gene and protein levels and the pro-coagulation activity were increased after incubating human umbilical vein endothelial cells with vape containing 18 mg/mL nicotine [27]. The inflammation that occurred in vaping groups increased the levels of IL-6, which consequently elevated platelet count and increased the tendency to blood coagulation and thrombosis. In addition, the presence of tissue factor leads to the acceleration in factor VIII conversion to its active form which will convert FX to its active form sequentially [28–30]. However, it has been approved by Cimmino et al, that VD can decrease tissue factor expression and atherosclerotic risk by modulating the nuclear factor kappa B in pre-incubated cells with VD. Although not all VD doses have the ability to decrease the activity of nuclear factor kappa B, the low dose VD of 1000 IU caused a reverse effect by increasing its activity in ulcerative colitis patients [31, 32]. That result clearly elucidates that 1000 IU of VD daily will cause an elevation in inflammatory biomarkers and increase the risk of thrombogenesis during stress conditions and inflammation. As the dose of VD increases, the activity of nuclear factor kappa B will be diminished in a dose-dependent manner during inflammation or stress conditions [33]. Those results explain the effect of 1000 IU and 50,000 IU on increasing and decreasing the inflammatory marker IL-6 and blood coagulation predictor levels in vape+50,000 IU VD and vape+1000 IU VD groups, respectively. In addition, the vape+1000 IU VD group had the highest creatine levels among all the groups with evidence of glomerulonephritis, which affected IL-6 clearance and led to its accumulation in the body.

The immunomodulatory action of VD is dose-dependent and changes according to physiological circumstances. According to Bock et al., giving healthy people 140,000 IU VD once a month markedly increased regulatory T cell activity, suggesting that large dosages of VD stimulate the immune system [34]. In a similar vein, Bader et al. showed that high-dose VD (50,000 IU) caused a cytokine storm and increased IL-6 levels in healthy individuals [35]. In contrast, under inflammatory or stress conditions, VD has been shown to reduce IL-6 levels in a dose-dependent manner [36].

D-dimer and IL-6 levels were shown to positively correlate in vaping-exposed groups, which is in line with research showing that fibrin breakdown products like D-dimer might increase vascular inflammation and IL-6 release [37]. Additionally, it has been shown that during inflammatory conditions, especially in lung infections as COVID-19, there is an inverse link between VD levels and D-dimer concentration [38]. The differences in D-dimer levels across groups might be explained by these interactions. The observed alterations in IL-6, D-dimer, TM, FX, ALT, and creatinine were probably caused by other components of the vape aerosol rather than nicotine alone, as evidenced by the notable lack of a significant link between nicotine and other measured parameters according to Pearson correlation analysis.

Regarding the histopathological findings, the necrotic results reported in the vape group lung tissue were consistent with an in vitro investigation conducted by Chastain Anderson et al, who discovered that vaping caused cell necrosis. Since TM is found in the alveolar epithelial cells, the drop in TM level that occurred after three months of exposure in the vape group was caused to necrotic pneumonia, which emerged after the histological examination as authorized by Boehme et al [39]. The Boehme et al. research discovered that TM levels increased dramatically before the start of cell injury, but the release was lost once cell damage occurred. These findings explain why the vape group had higher levels of IL-6 and lower levels of TM after one and three months, respectively [40]. The bacterial pneumonia that emerged in the vape+50,000 IU VD group may be attributed to the absence of a VD receptor, which impacts the innate defense against bacterial infections in rodents and is confined to primates [41].

As the hepatic regeneration capacity in Wistar rats reaches its maximum ability between 9 to 24 weeks of age [41]. This could strongly explain the histopathological examination of the liver, which showed no significant changes in the hepatic tissues after three months of exposure.

The glomerular histopathology examination showed that there was immune-related kidney injury. This damage was produced by inflammation and activation of the NF-kB pathway, which led to an increase in IL-6 levels. This pathway activation increases the synthesis of proinflammatory cytokines present in the nephritic glomeruli, resulting in glomerulonephritis in rats [42].

## Conclusion

In conclusion, our study shows that short-term nicotine vape exposure increases coagulability indicators and inflammatory cytokines in female rats, but longer exposure may cause physiological adaptation. High-dose VD supplementation (50,000 IU/week) protected vaping-exposed rats against these changes, whereas low-dose VD (1,000 IU/day) showed moderate efficacy. Notably, high-dose VD in non-vaping rats induced a proinflammatory response. These data indicate that VD may reduce vaping-induced coagulopathy and inflammation while also potentially aiding in nicotine detoxification. While based on an animal model, the findings emphasize the potential therapeutic benefit of VD in vape users.

## Declaration

## Acknowledgment

The authors thank X-vape shop, Smart lab, and Al-Hikma lab.

## Data availability

All relevant data can be found within the manuscript and in the supporting file.

## Author contributions

Conceptualization: Muna Barakat and Mahmoud Abu-Samak.

Methodology: Lujain F. Zghari, Aman M. Hammad.

Writing - original draft: Aman M. Hammad.

Writing – review and editing: Muna Barakat.

---

**Figure S4.** Summary of mean plasma (A) D-dimer concentrations (ng/mL), (B) TM concentrations (ng/mL), and (C) FX concentrations (µg/mL) among all the groups at baseline, one month, and three months.

**Figure S5.** (A) Summary of mean plasma ALT concentrations (U/L), and (B) Creatinine concentrations (µmol/L) among all the groups at baseline, one month, and three months.

**Figure S6.** Histopathological appearance of liver tissue from all the groups under light microscopy.

**Figure S7.** Histopathological appearance of kidney tissue from all the groups under light microscopy.

**Figure S8.** Histopathological appearance of lung tissue from all the groups under light microscopy.

